# AuCoMe: inferring and comparing metabolisms across heterogeneous sets of annotated genomes

**DOI:** 10.1101/2022.06.14.496215

**Authors:** Arnaud Belcour, Jeanne Got, Méziane Aite, Ludovic Delage, Jonas Collen, Clémence Frioux, Catherine Leblanc, Simon M. Dittami, Samuel Blanquart, Gabriel V. Markov, Anne Siegel

**Author notes:** Corresponding authors: Arnaud Belcour and Anne Siegel;. These authors contributed equally to this work (Arnaud Belcour, Jeanne Got, Méziane Aite). Co-last authors (Gabriel V. Markov, Anne Siegel).

## Abstract

Comparative analysis of Genome-Scale Metabolic Networks (GSMNs) may yield important information on the biology, evolution, and adaptation of species. However, it is impeded by the high heterogeneity of the quality and completeness of structural and functional genome annotations, which may bias the results of such comparisons. To address this issue, we developed AuCoMe – a pipeline to automatically reconstruct homogeneous GSMNs from a heterogeneous set of annotated genomes without discarding available manual annotations. We tested AuCoMe with three datasets, one bacterial, one fungal, and one algal, and demonstrated that it successfully reduces technical biases while capturing the metabolic specificities of each organism. Our results also point out shared metabolic traits and divergence points among evolutionarily distant species, such as algae, underlining the potential of AuCoMe to accelerate the broad exploration of metabolic evolution across the tree of life.

## Introduction

The comparison of genomic data gave rise to today’s view of the three domains of life: bacteria, archaea, and eukaryotes, the latter being divided into five putative supergroups, opisthokonts, amoebozoans, archaeplastids (including green and red algae lineages), Stramenopiles-Alveolata-Rhizaria (including brown algae) and Excavata, and additional uncertain lineages (Burki *et al*, 2020). The evolution of the organisms within these lineages is linked to their ability to adapt to their environment and, therefore, to the plasticity of their metabolic responses.

In this context, the comparison and analysis of Genome-Scale Metabolic Networks (GSMNs) constitute a powerful approach (Gu *et al*, 2019). The number of sequences available in public databases is continuously rising, as illustrated by the GenBank database, which grew by 74.30% for Whole Genome Shotgun data in 2019 compared to 2018 (Sayers *et al*, 2019), and GSMN reconstruction is theoretically possible for any genome and has already been used to explore evolutionary questions. In Schulz and Almaas (2020), metabolic relationships were explored in 975 organisms from the three domains of life, showing that these domains were well-separated in terms of metabolic distance despite some organisms being misplaced. Using GSMN reconstruction in bacteria, metabolic and phylogenetic distances between *Escherichia coli* and *Shigella* strains could be explained by the parasitic lifestyle of the latter (Vieira *et al*, 2011). Another GSMN-based study of 301 genomes from the human gut microbiota identified marginal metabolic differences at the family level but significant metabolic differences between closely related species (Bauer *et al*, 2015). Analysis of fungal GSMNs additionally demonstrated correlation between metabolic distances and the phylogeny of *Penicillium* species, even if no connection was found between the metabolic distances and the species habitat (Prigent *et al*, 2018). In brown algae, the GSMNs of *Saccharina japonica* and *Cladosiphon okamuranus* (Nègre *et al*, 2019) were compared to the GSMN of *Ectocarpus siliculosus* (from Prigent et al (2014)): a thorough analysis of the differences between their genomic contents revealed that heterogeneity of genome annotations may have a stronger impact on GSMNs than genuine biological differences.

For most GSMN analyses, some limitations still need to be addressed, as recently reviewed (Bernstein *et al*, 2021). When comparing different GSMNs, two main biases concern the variable quality of genome annotations and the multitude of reconstruction approaches. Indeed, a variety of methods exists to perform structural (gene structure prediction) and functional (association of functions to genes) annotation steps (Yandell and Ence, 2012) and the method choice has previously been shown to have direct effects on the reconstructed GSMNs (Karimi *et al*, 2021). Similarly, numerous methods for GSMN reconstruction have been developed, e.g. Pathway Tools (Karp *et al*, 2019), RAVEN (Wang *et al*, 2018), merlin (Dias *et al*, 2015a), KBase (Arkin *et al*, 2018), Model SEED (Devoid *et al*, 2013), AuReMe (Aite *et al*, 2018), AutoKEGGRec (Karlsen *et al*, 2018), CarVeMe (Machado *et al*, 2018), and gapseq (Zimmermann *et al*, 2021a). They rely on one or several metabolic databases such as MetaCyc (Caspi *et al*, 2020), KEGG (Kanehisa and Goto, 2000; Kanehisa et al, 2017), Model SEED (Seaver *et al*, 2021) or BiGG (King *et al*, 2016). Despite efforts in the direction of database reconciliation (Moretti *et al*, 2021), such heterogeneity of metabolic databases still requires time-consuming matching of their respective identifiers and may thus impede the comparison of GSMNs generated.

One strategy to resolve the issue of GSMN comparison is to work directly on GSMNs. A first method is the *reconstruction annotation jamboree* (Thiele and Palsson, 2010), a community effort to curate pathway discrepancies by examining reactions, Gene-Protein-Reaction (GPR) associations, and metabolites in GSMNs in order to create a consensus GSMN for an organism. This is relevant for organisms for which multiple GSMNs exist, in order to establish a reference one. This strategy was successfully applied to *Salmonella Typhimurium LT2* (Thiele *et al*, 2011) as well as *Saccharomyces cerevisiae* (Herrgård *et al*, 2008), and was later extended to multiple organisms to create a panmetabolism of 33 fungi from the Dikarya subkingdom (Correia and Mahadevan, 2020). Although platforms now facilitate such community efforts (Cottret *et al*, 2018), these methods are costly in terms of time and manpower involved. Another method was proposed by Oberhardt et al (2011) to resolve the comparison of GSMNs by reconciling their GPR associations using reciprocal gene pairs.

A second strategy to resolve GSMN comparison issues is to adapt the GSMN reconstruction method used to tackle these issues. This strategy aims at reducing annotation biases through the reconstruction of GSMNs from homogeneously annotated genomes using the same method and database, possibly followed by the propagation of annotations with sequence alignments (Vieira *et al*, 2011; Prigent et al, 2018). This strategy was pushed forward and automatized in the tool CoReCo, which enabled the reconstruction of gap-less metabolic networks from several non-annotated genomes (Pitkänen *et al*, 2014; Castillo et al, 2016). The main limitation of such approaches is that the re-annotation of the genomes supplants the previous genome annotation.

Annotations of genomes in databases also reflect the expertise of scientists. Such valuable information, possibly originating from manual curation, would be lost during a systematic re-annotation step. For a reliable interpretation of data, expert annotations therefore ought to be preserved while automatically inferring metabolic networks from any type of genomic resource. In this article, we introduce a new method, *AuCoMe* (Automated Comparison of Metabolism) that creates a set of homogenized GSMNs from heterogeneously-annotated genomes. This enables the less biased functional comparison of the networks and the determination of metabolic distances using the presence/absence of reactions. Our objective was to develop an efficient, fast, and robust approach, which does not depend on the quality of the initial annotations and is able to aggregate heterogeneous information. AuCoMe combines metabolic network reconstruction, propagation, and verification of annotations. The method automatizes the strategy of transferring information from the annotations of the genomes and complements this information transfer with local searches of missing structural annotations. AuCoMe was applied to three heterogeneous datasets composed of fungal, algal, and bacterial genomes, respectively. Our results demonstrate that AuCoMe succeeds at propagating missing reactions to degraded metabolic networks while capturing the metabolic specificities of organisms despite profound differences in the quality of genome annotations. Finally, our results illustrate that the analysis of the panmetabolism provides a knowledge base for the comparison of metabolisms between different organisms, especially for the identification of shared and diverging metabolic traits.

## Results

### A tool for homogenizing metabolism inference

AuCoMe is a python package that aims to build homogeneous metabolic networks and panmetabolisms starting from genomes with heterogeneous functional and structural annotations. AuCoMe propagates annotation information among organisms through a four-step pipeline (Fig.1). As a first step, draft metabolic networks are automatically inferred from the original annotations of genomes using Pathway Tools (Fig.1A). By dsefault, only reactions supported by gene associations are kept in the draft metabolic networks (see Methods). The resulting GSMNs and their proteomes are then subjected to comparative genomic analyses. During this process, GPR associations are propagated across GSMNs according to orthology relations established using OrthoFinder (Fig.1B). A robustness filter (see Methods) then selects the robust GPR relationships among all propagated associations. A third step consists in checking for the presence of additional GPR associations by finding missing structural annotations in all genomes (Fig.1C). Finally, a fourth step adds spontaneous reactions associated with MetaCyc metabolic pathways that were completed by the two previous steps (Fig.1D).

**Figure 1:**
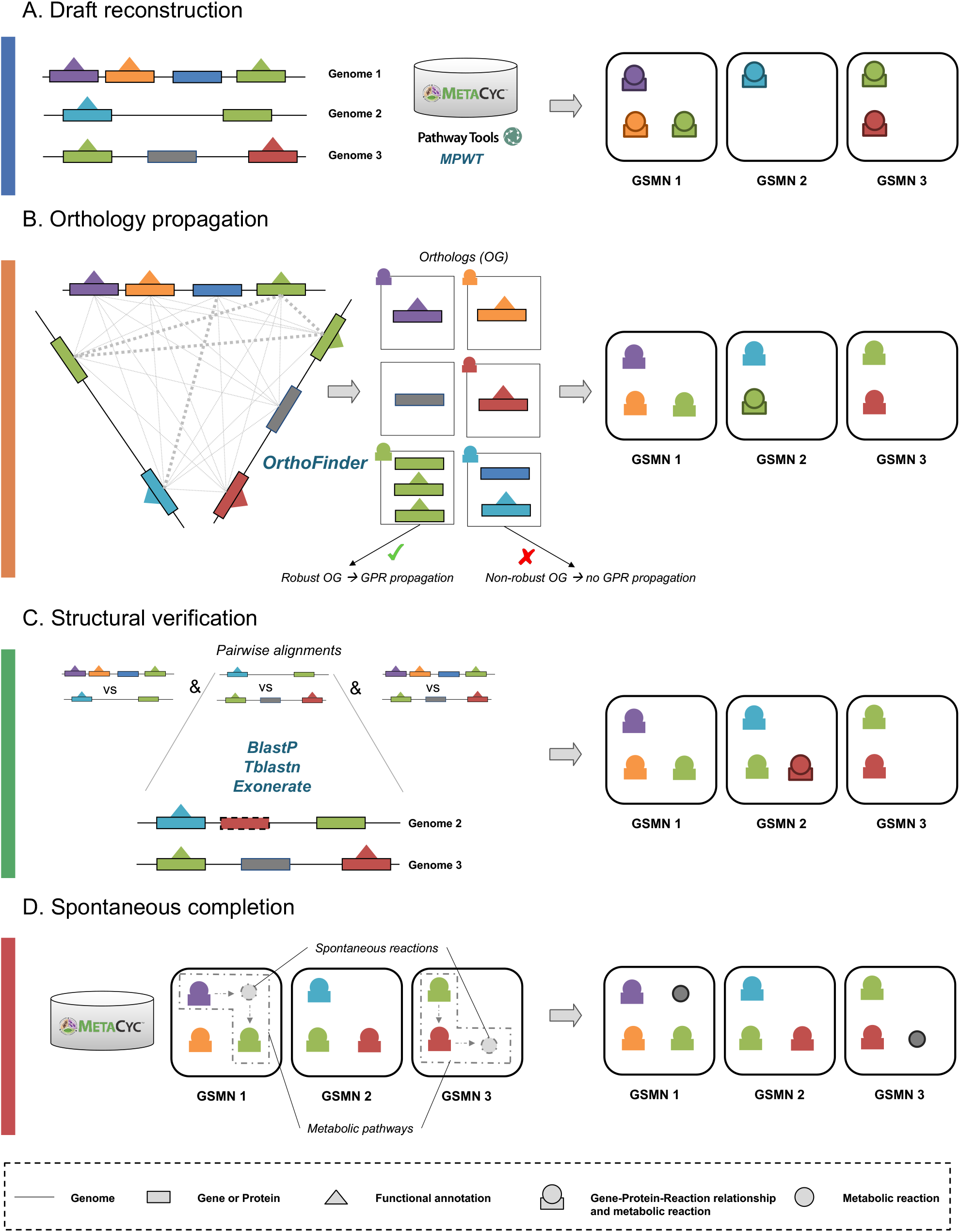
Reconstruction and homogenization of metabolisms with AuCoMe. Starting from a dataset of partially structurally-and functionally-annotated genomes, the AuCoMe pipeline performs the following four steps. **A. Draft reconstruction**. The reconstruction of draft genome-scale metabolic networks (GSMNs) is performed using Pathway Tools in a parallel implementation. **B. Orthology propagation**. OrthoFinder predicts orthologs by aligning protein sequences of all genomes. The robustness of orthology relationships is evaluated (see Methods), and GPRs of robust orthologs are propagated. **C. Structural verification**. The absence of a GPR in genomes is verified through pairwise alignments of the GPR-associated sequence to all genomes where it is missing. If the GPR-associated sequence is identified in other genomes, the gene is annotated, and the GPR is propagated. **D. Spontaneous completion**. Missing spontaneous reactions enabling the completion of metabolic pathways are added to the GSMNs. GSMN: Genome-scale metabolic network. OG: orthologs. GPR: Gene-protein-reaction relationship.

The AuCoMe pipeline was tested on three datasets composed of genomes that offer different levels of phylogenetic diversity. The *bacterial dataset* includes 29 genomes belonging to different species of *Escherichia* and closely related *Shigella*, the *fungal dataset* (74 fungal genomes and 3 outgroup genomes) covers a range of different phyla within this kingdom, and finally the *algal dataset* (36 algal genomes and 4 outgroup genomes) exhibits the highest phylogenetic diversity including eukaryotes from the supergroups SAR (Stramenopiles, Alveolata and Rhizaria), Haptophyta, Cryptophyta, and Archaeplastidia. For all species included in the three datasets, both annotated genomes and proteomes were publicly available (see Appendix Tables S1, S2, S3).

Run times of AuCoMe on a cluster were 7 hours (10 CPUs), 25 hours (40 CPUs), and 45 hours (40 CPUs) for the bacterial, fungal, and algal datasets, respectively. Details for individual steps are reported in Appendix, section 2.

### AuCoMe homogenizes the content of metabolic network collections

In Fig. 2, we compared the number of reactions inferred during the different steps of the AuCoMe pipeline.

**Figure 2:**
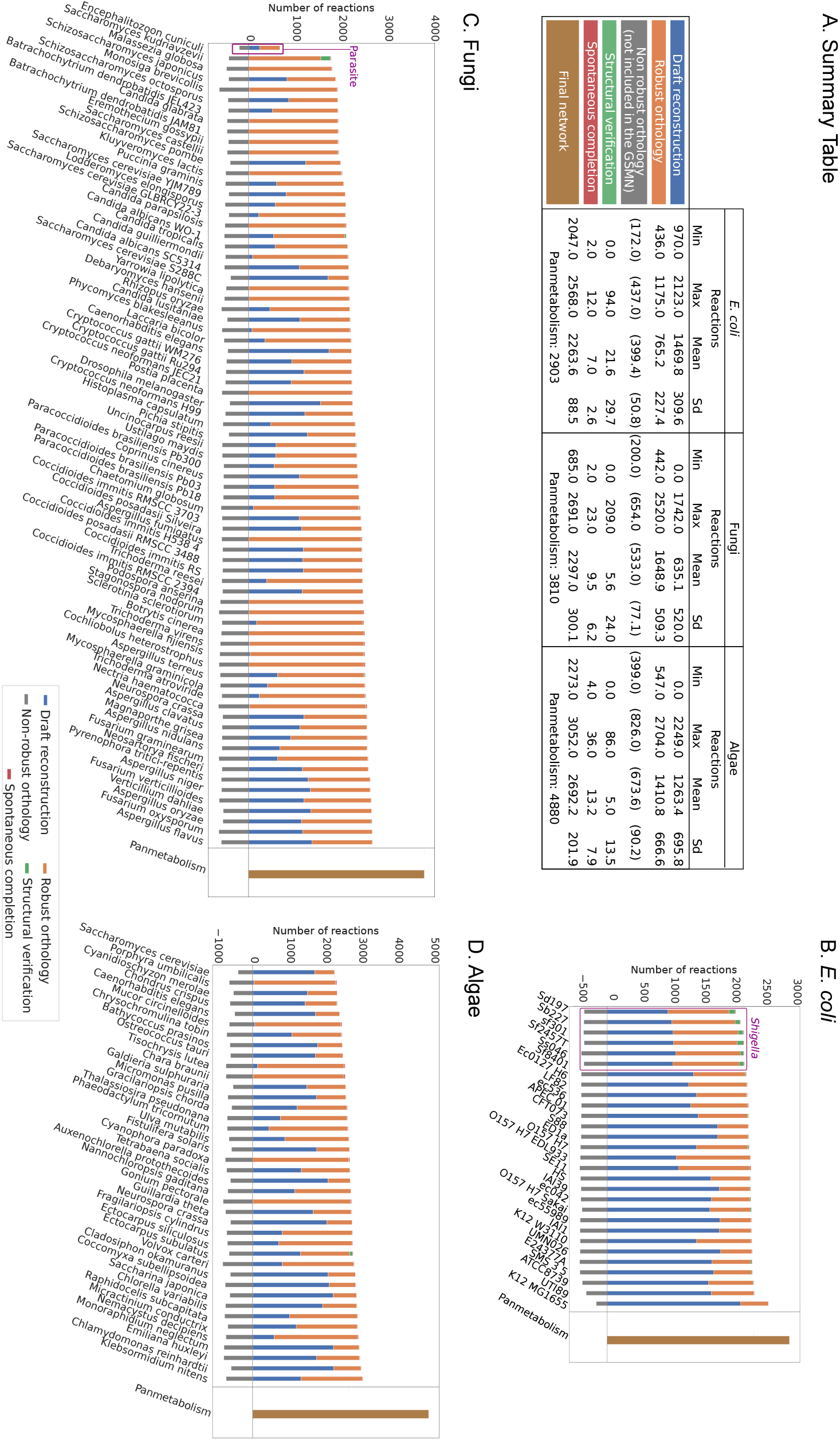
Application of the AuCoMe pipeline to the *bacterial, fungal* and *algal* datasets of genomes. The figure depicts the number of reactions identified for each species at each step of the Au-CoMe pipeline: reactions recovered by the *draft reconstruction* step (blue), unreliable reactions predicted by orthology propagation and removed by the filter (gray), robust reactions predicted by *orthology propagation* that passed the filter (orange), additional reactions predicted by the *structural verification* step (green), and *spontaneous completion* (red). The final metabolic networks encompass all these reactions except the non-reliable ones. The panmetabolism (all the reactions occurring in any of the organisms after the final step of AuCoMe) is presented in brown in B, C and D.

After the *draft reconstruction* step, draft GSMNs from the three datasets exhibit a highly heterogeneous range of reactions (Fig. 2 A and blue bars in Fig. 2 B, C, D). This is especially visible for the fungal dataset, demonstrating the variability of the original genomic annotations. In this dataset, no reactions were inferred from annotations in seven species, and 12 draft GSMNs contained less than ten reactions. For the latter, their respective genome annotations included no EC number, and eleven genomes did not include any GO term. As Pathway-Tools relies mainly on these two annotations to infer reactions, this absence impedes the reconstruction process. Similar observations were also made, although to a lesser extent, for the algal genome dataset, with seven genomes having more than 2,000 reactions and seven genomes less than 500 reactions. At this step, high heterogeneity in the number of reactions can be attributed mainly to differences in the quality and quantity of the functional annotations provided, precluding biologically meaningful comparisons of the GSMNs obtained at the draft reconstruction step.

After the *orthology propagation* step, we observed an homogenization of the number of reactions in the datasets (orange bars in Fig. 2). GSMNs with few reactions after the draft reconstruction recovered more reactions during the orthology propagation step compared to GSMNs with thousands of reactions. This observation is supported by the negative correlation between the number of reactions added at the draft reconstruction step and at the orthology propagation step (Spearman’s rs -0.82 p¡0.001). The fungal dataset exhibits an outlier at this step; the GSMN of *Encephalitozoon cuniculi* contained only 681 reactions compared to the thousand of reactions in the other fungal GSMNs. This is consistent with the fact that this species is a microsporidian parasite with a strong genome and gene compaction (Grisdale *et al*, 2013). In all datasets, among the reactions propagated byorthology, a few hundred were removed because they did not fulfill the robustness score criterion (see methods).

Compared to the orthology propagation, the *structural verification* step had a smaller impact on the size of the final networks (green bars in Fig. 2). Ninety-five percent of the GSMNs received less than 28 reactions during this step, and the maximum was 209. In the bacterial dataset, the six *Shigella* received an average of 76.2 reactions which is more than ten times the average of the other strains (7.4). We manually examined these differences and found that the majority of genes in the GPR associations added at this step in the *Shigella*’s GSMNs corresponded to pseudogenes. For the fungal dataset, AuCoMe added 209 reactions for the species *Saccharomyces kudriavzevii*. These reactions were associated with 192 sequences recovered during the structural step. For all of these sequences, we found corresponding transcripts in a previously published transcriptome dataset (Blevins *et al*, 2021). As for the algal dataset, 86 reactions were added for *Ectocarpus subulatus*. Likewise, we validated the presence of 59 out of 65 genes (83 out of 86 reactions) by associating them with existing transcripts. The remaining six genes (three reactions) corresponded to valid plastid sequences that had remained in the nuclear genome assembly. These genes lacked transcription data, because plastid mRNA lacks the PolyA tail used to prepare most RNAseq libraries. In both outlier cases, the structural completion step was, therefore, able to recover sequences that are translated into mRNA and thus likely to correspond to functional genes.

Lastly, the *spontaneous completion* step added spontaneous reactions to each metabolic network if these reactions complete BioCyc pathways (red bars in Fig. 2). For the fungal dataset, this step added between two and 23 spontaneous reactions, leading to two to 27 additional MetaCyc pathways that achieved a completion rate greater than 80%. As for the algae, the same step added between 4 and 36 spontaneous reactions yielding two to 31 additional pathways that reached a completion rate greater than 80%. Generally, we observed that the fewer reactions were inferred at the draft reconstruction step, the more spontaneous reactions were added to complete pathways (Pearson R = -0.83 and -0.84 for the fungal and algal datasets, respectively). This can be explained by the fact that the only other possibility to introduce spontaneous reactions in the GSMNs is the draft reconstruction step because Pathway Tools infers GSMN with both enzymatic and spontaneous reactions.

When looking at the size of the final networks, we observed in the bacterial dataset (Fig. 2B) that the networks of *Shigella* strains comprised fewer reactions than the rest (average of 2,148 reactions vs. 2,294, Wilcoxon rank-sum test W = 138, P = 2e-4). This is consistent with the results of Vieira *et al* (2011) (average of reactions of 1,437 for *Shigella* strains compared to 1,504 for non-*Shigella* strains). On the other hand, *E. coli* K–12 MG1655 stood out with 2,568 reactions compared to the range of reactions between 2,047 and 2,342 for the other strains. An explanation is that this strain is clearly the best annotated of all tested strains, and the corresponding reference genome in the databases has been extensively curated. For this strain, 172 reactions were removed by the robustness filter, which is less than the 333-447 reactions removed for the other strains. The difference between these two numbers can be explained because reactions propagated from *E. coli* K–12 MG1655 to the other strains were frequently supported by only one gene predicted at the draft reconstruction step, and were removed after the orthology propagation (see Methods). Overall, in the three datasets, the final GSMNs were of similar size after applying AuCoMe regardless of the quantity and quality of their corresponding genome annotations.

### Validation of filtering steps and final GPR associations

To validate the above results, two independent verification procedures were performed, the first one at the level of the orthology propagation step in order to evaluate the efficiency of the propagation filters, and the second one applying manual examination of the final networks for the algal dataset.

For the first verification procedure, we extracted the EC numbers of all reactions of the fungal and the algal dataset GSMNs for which GPR associations were only predicted by orthology propagation. For each EC number, we then extracted the associated protein sequence and used DeepEC (Ryu *et al*, 2019) to infer EC numbers for these proteins and compared them (if any) to the EC number(s) linked to the reaction by the pipeline. An enrichment of sequences confirmed by DeepEC is observed in robust GPR associations compared to those discarded by the filter: 24% vs. 4.9 % in the fungal dataset and 13% vs. 1.4% in the algal dataset (see Appendix Fig S3). This confirms that the robustness filter removed predominantly poorly supported reactions.

The second evaluation of the reliability of the reconstruction process was performed on the final algal dataset. We manually examined 100 random GPR associations across the metabolic networks generated by AuCoMe: 50 reactions that were predicted to be present and 50 reactions that were predicted to be absent (see methods). Not counting spontaneous reactions (seven among the randomly selected reactions), manual annotations and automatic predictions corresponded in 86% of all cases (42/49) for the reactions predicted to be present and in 91% (40/44) for the reactions predicted to be absent (see Appendix Tables S5 and S6). These data underline the robustness of the AuCoMe pipeline.

### Validation of the orthology propagation and structural verification steps

Thirty-two datasets were formed, each containing the 29 bacterial *E. coli* and *Shigella* strains studied in Vieira et al (2011), among them a replicate of the *E. coli* K–12 MG1655 genome being degraded to a variable extent, in its functional and/or structural annotations (see Methods and Appendix Table S4). The manually-curated EcoCyc database (Karp *et al*, 2018) was used to check the reliability of the GSMN reconstructed for each corresponding degraded genome. For each of the 32 datasets, F-measures were computed at each AuCoMe step according to comparisons of the reconstructed GSMN with the gold-standard EcoCyc database (see Methods). The F-measure stands for the harmonic mean of the AuCoMe precision and recall, with values close to 1 indicating high precision and recall *i*.*e*. low rates of false positives and false negatives, respectively. The maximal theoretical value of the F-measure in our analyses was obtained by computing the maximal F-measure that can be reached when comparing to EcoCyc. To achieve this, we computed the number of reactions contained in EcoCyc but not in the panmetabolism of dataset 0, which is the union of GSMNs created by AuCoMe from all the genomes of dataset 0, including the non-degraded annotations of the *E. coli* K–12 MG1655 genome. Among the reactions of EcoCyc, 1,019 reactions could not be retrieved in this panmetabolism. This number of reactions corresponded to the false negatives, the other reactions of EcoCyc corresponded to the true positives and using these measures we computed the maximal theoretical F-measure of 0.79. A first analysis was performed on the non-degraded dataset 0, and after running the AuCoMe process, the F-measure between the GSMN of the *E. coli* K–12 MG1655 genome in dataset 0 and EcoCyc was 0.67. Therefore we consider 0.67 as the reference F-measure for this experiment.

Fig. 3 A illustrates the number of reactions predicted by the steps of AuCoMe for the *E. coli* K–12 MG1655 GMSN in each of the 32 synthetic bacterial datasets to assess the importance of each step in the homogenization of the GSMN sizes. Fig. 3 B represents the F-measure for the corresponding dataset. As expected, increasing degradation led to the recovery of fewer GPR associations and decreased F-measures in the draft GSMN (blue bars and blue circles). When all annotations were removed (dataset 10, 21 and 31) both the number of reactions and F-measure were closed to 0. When only functional annotations were degraded in the *E. coli* K–12 MG1655 genome (dataset labeled 1 to 10), the orthology propagation step enabled the recovery of a large number of reactions, nearly up to the size of the GSMN associated with the non-degraded genome. This also resulted in increasing F-measures (orange bars and orange triangles). In datasets where only structural annotations of the *E. coli* K–12 MG1655 genome were altered (datasets labeled 22 to 31), the structural verification step recovered a part of the GPR associations and a F-measure close to the reference F-measure of 0.67 (green bars and green crosses). Finally, when both structural and functional annotations were degraded (datasets 11 to 21), the combination of the second and third AuCoMe steps recovered the discarded reactions. Notably, even when 100% of the *E. coli* K–12 MG1655 functional and structural annotations are degraded, the information from the other 28 non-altered genomes enabled the recovery of 2,244 reactions (Fig. 3 A, dataset 31) and a F-measure of 0.60.

**Figure 3:**
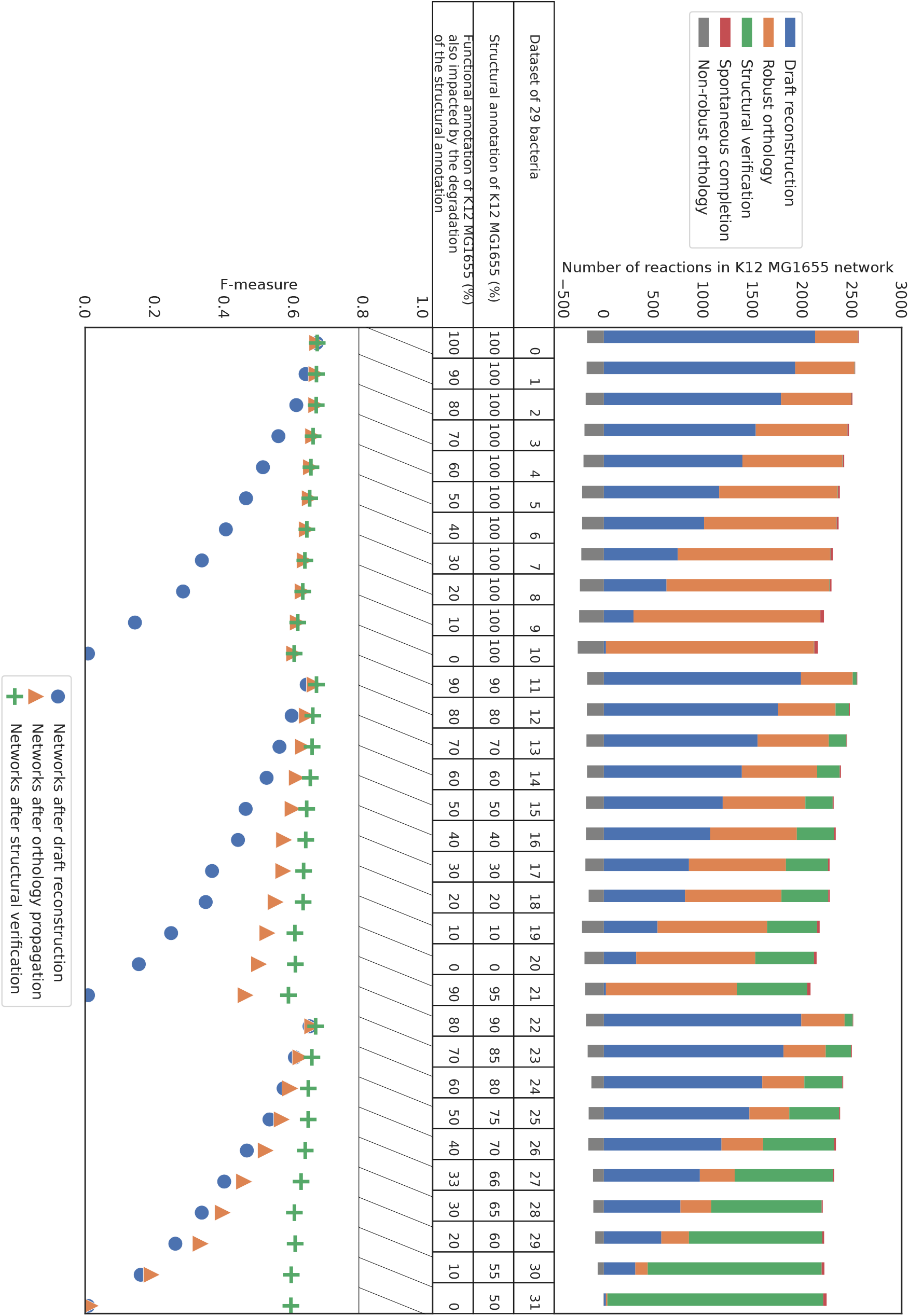
(A) Number of reactions in *E. coli* K–12 MG1655 degraded networks after application of AuCoMe to 32 synthetic bacterial datasets. Each dataset consists of the genome of *E. coli* K–12 MG1655 to which degradation of the functional and/or structural annotations was applied, together with 28 bacterial genomes. Each vertical bar corresponds to the result on the *E. coli* K–12 MG1655 within a synthetic dataset, with the percentages of degraded annotations indicated below. The dataset labelled 0 was not subject to degradation of the *E. coli* K–12 MG1655 annotations. Three types of degradation were performed: functional annotation degradation only (left side, datasets labelled 1 to 10), structural annotation degradation only (right side, datasets labelled 22 to 31) and both degradation types (middle, dataset labelled 11 to 21). The colored bars depict the number of reactions added to the degraded network at the different steps of the method (the blue, orange, green, grey, and red color legends are as described in the figure 2). The table shown as axis indicates the dataset number and the percentage of functional or structural annotation impacted by the degradation for the corresponding column in both subfigures. **(B) F-measures after comparison of the GSMNs recovered for each *E. coli* K–12 MG1655 genome replicate with a gold-standard network**. Reactions inferred by each AuCoMe step for each replicate were compared to the gold-standard EcoCyc GSMN, allowing for the computation of F-measures. F-measures obtained after the draft reconstruction step, the orthology propagation step, or the structural verification step are shown as blue circles, orange triangles, and green crosses, respectively. The hashed rectangle from F-measure 0.79 to 1 highlights the values of F-measure that are unreachable because 1019 reactions in EcoCyc were not present in the panmetabolism of the 29 non-degraded bacteria.

Altogether, these results demonstrate that, by taking advantage of the annotations present in the other genomes of the considered dataset, AuCoMe builds GSMNs with reactions even for genomes completely missing functional and structural annotations.

### Biological validation on the Calvin pathway

The Calvin cycle is a pathway of biochemical reactions present in photosynthetic organisms to fix carbon from *CO*_2_ into three-carbon sugars. In the MetaCyc v.23.5 database (Karp *et al*, 2019), the Calvin cycle (id CALVIN-PWY) is composed of 13 reactions. In Fig. 4, the different reactions (rows) predicted for each genome of the algal dataset (columns) are depicted together with the step of AuCoMe at which the prediction occurred. In this example, all three main AuCoMe steps are required to obtain a homogeneous view of this pathway in all organisms. The draft reconstruction (blue) and the orthology propagation (orange) steps provide most of the reactions. Interestingly, the pathway was predicted to be complete for three species *G. pectorale, C. braunii* and *C. paradoxa* although their genomes did not contain any annotation related to this pathway. The structural verification step added one reaction (RIBULP3EPIM-RXN) for *P. umbilicalis* (green square in Fig. 4).

**Figure 4:**
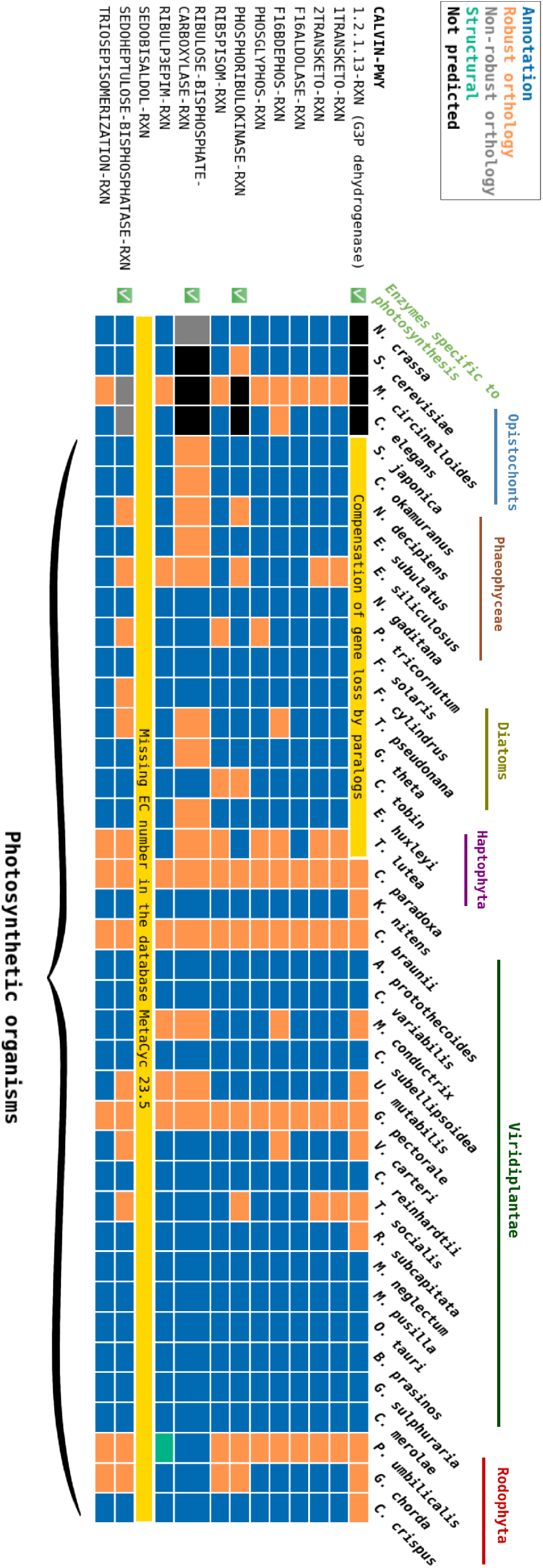
AuCoMe results on the Calvin cycle pathway in the algal dataset. AuCoMe was applied to the dataset of 36 algae and 4 outgroup species (columns). Each row represents a MetaCyc reaction of the pathway, the table shows whether it is predicted by AuCoMe: blue - draft reconstruction, orange - robust reactions predicted by orthology propagation that passed the filter, green - structural verification, and gray - non-robust reactions predicted by orthology propagation and removed by the filter, black - not predicted, yellow - manually added because the MetaCyc database 23.5 does not contain a reference gene-reaction association for this reaction).

After the reconstruction, the genomes can be clustered into three groups with respect to the Calvin Cycle. The first group, containing the green algal (Viridiplantae), red algal (Rhodophyta), and *C. paradoxa* genomes, was predicted to contain 12 reactions of the pathway. The second group consisted of the four outgroup organisms that do not possess the 1.2.1.13-RXN and RIBULOSE-BISPHOSPHATE-CARBOXYLASE-RXN reactions. The last group contained brown algae, diatoms, haptophytes, *G. theta* and *N. gaditana* that lack the 1.2.1.13-RXN reaction associated with the enzyme Glyceraldehyde-3-phosphate dehydrogenase (GAPDH).

None of the 40 GSMNs contained the reaction SEDOBISALDOL-RXN. In the MetaCyc v23.5 database, which was used for the experiment, this reaction was associated with the EC number 4.1.2.X, whereas it was associated with the EC number 4.1.2.13 in the KEGG (Kanehisa *et al*, 2008) and Brenda (Schomburg *et al*, 2017) databases. This prevented the draft reconstruction tool (Pathway Tools v23.5) from associating genes annotated with EC 4.1.2.13 to this reaction. We therefore manually added the corresponding GPR associations, indicated in yellow in Fig. 4.

Regarding the outgroup organisms, the robustness criterion applied during the orthology propagation step removed a weak GPR association with the reaction RIBULOSE-BISPHOSPHATE-CARBOXYLASE-RXN for *N. crassa*. This weak association was previously reported and explained by the sharing of a SET domain by Rubisco and other lysine protein methyltransferases (Yeates, 2002). Similarly, the protein annotated as phosphoribulokinase in *S. cerevisiae* and *N. crassa* is an ATP-dependent kinase reported to exhibit a weak similarity with the phosphoribulokinase and other P-loop containing kinases (Gueguen-Chaignon *et al*, 2008). Finally, the annotation of a sedoheptulose bisphosphate in *S. cerevisiae* and *N. crassa* has previously been reported in Ascomycete fungi, where it is assumed to be unrelated to photosynthesis (Teich *et al*, 2007).

In the last group, the missing GAPDH (1.2.1.13-RXN) reaction could be explained by the fact that AuCoMe searches for orthologs during the orthology propagation step using OrthoFinder. The GAPDH reactions have been propagated to green algae, red algae and *C. paradoxa* because they share a common ortholog associated with this reaction, the plastid gene *GapA*, whereas this gene has not been found in diatoms and brown algae. This could be explained by the fact that a duplicated cytosolic GAPDH can functionally replace this gene (Liaud *et al*, 1997). The presence of these genes was validated using sequence alignment with known cytosolic GAPDH and the reaction was manually added (yellow).

### Prediction of pigment pathways

We used AuCoMe to predict the capacity of five brown algae, *Cladosiphon okamuranus, Ectocarpus siliculosus, Ectocarpus subulatus, Nemacystus decipiens*, and *Saccharina japonica* to synthesize heme groups and then to catalyze three alternative pigment biosynthesis pathways in the MetaCyc database: phycoerythrobilin (PWY-5915), phycocyanobilin (PWY-5917), and phytochromobilin (PWY-7170) (Fig. 5). These pathways consist of four, three, and three reactions, respectively, and all have protoheme, the end-product of the heme synthesis pathway, as a starting point. They also share the same starting reaction (RXN-17523) that transforms heme into biliverdin in the presence of a reduced ferredoxin (Fig. 5).

**Figure 5:**
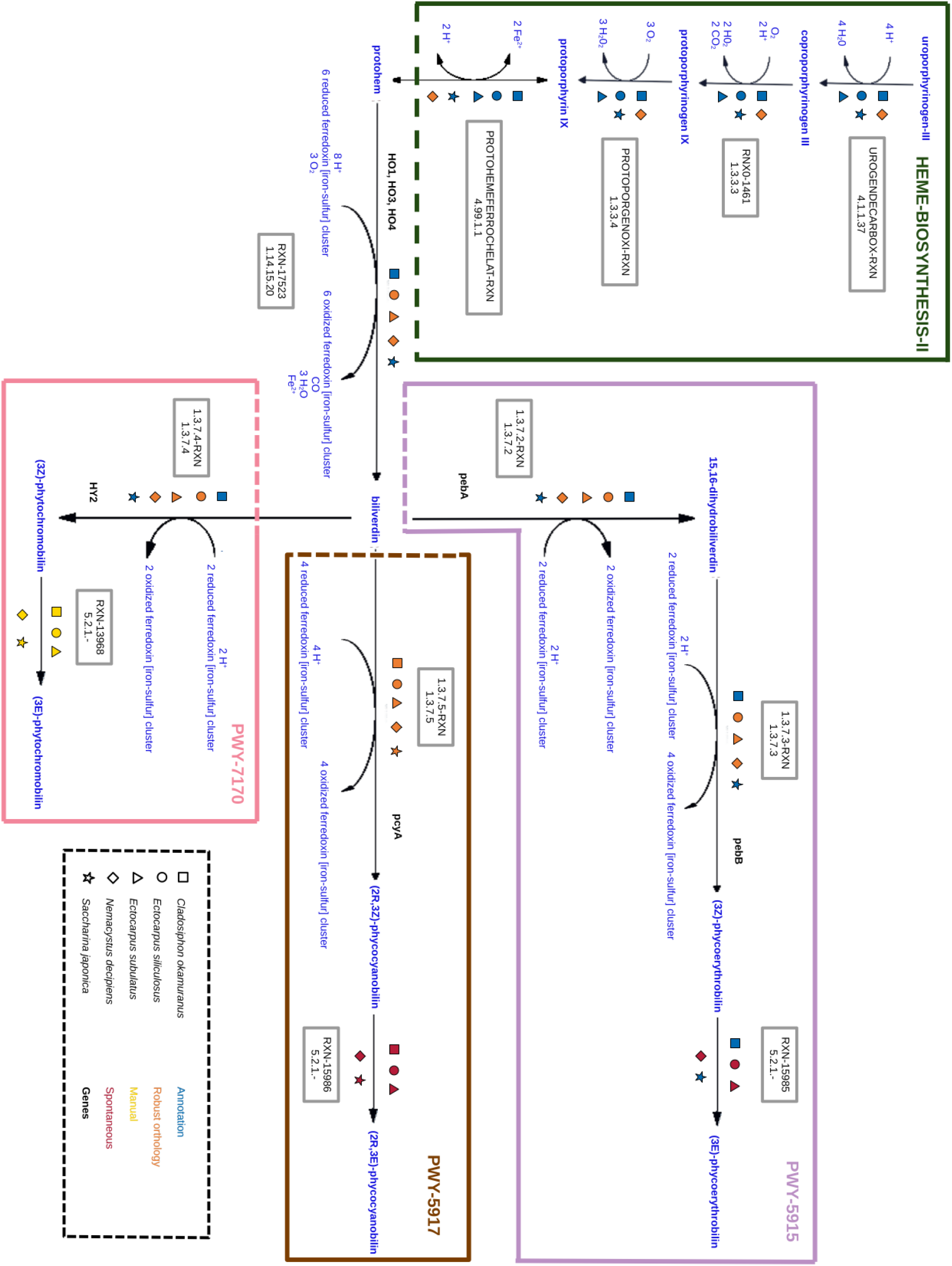
Prediction of pigment pathways in brown algae. Links between four MetaCyc pigment pathways are presented: heme b biosynthesis (HEME-BIOSYNTHESIS-II, dark green frame), phycoerythrobilin biosynthesis (PWY-5915, mauve frame), phycocyanobilin biosynthesis (PWY-5917, brown frame), and phy-tochromobilin biosynthesis (PWY-7170, pink frame). PWY-5915, PWY-5917, and PWY-7170 are three alternative biosynthetic pathways of bilin proteins. All three follow the heme B biosynthesis, and start with reaction RXN-17523. Here we detailed every AuCoMe step employed to complete the GSMNs of five brown algae, from the GPR associations found with draft reconstruction (blue), via GPR associations specifically found by orthology propagation, to GPR associations that passed the filter (orange). In the end, AuCoMe also added missing spontaneous reactions to GSMNs (see Materials & Methods). Furthermore, we chose to manually add a few reactions (gold) using the PADMet library (Aite *et al*, 2018). We focused on five brown algae: *C. okamuranus* (square), *E. siliculosus* (circle), *E. subulatus* (triangle), *N. decipiens* (diamond), and *S. japonica* (star). Genes are noted in bold black.

Regarding the heme biosynthesis pathway, the added value of AuCoMe was to predict GPR associations for *N. decipiens* - whose genome had few functional annotations - for each of the reactions of the pathway. The reaction RXN-17523 and all other enzymes of PWY-5915 were predicted to be present in *C. okamuranus* and *S. japonica* based on their genome annotations. These enzymes were also found in the three other algae by orthology propagation. The other reactions in PWY-5917 are 1.3.7.5-RXN and RXN-15986. AuCoMe added 1.3.7.5-RXN to all brown algal GSMNs based on orthology with several green algae. These added associations were considered robust by the software. RXN-15986 is a spontaneous reaction that was added by AuCoMe. In the same vein, in addition to RXN-17523, PWY-7170 comprises the reactions 1.3.7.4-RXN and RXN-13968. 1.3.7.4-RXN was predicted to be present in *S. japonica* based on annotations and predicted to be also present in other brown algae, based on orthology. RXN-13968 (Karp *et al*, 2019) was not considered spontaneous but had no associated enzymes in the MetaCyc database and hence was not found in any of the analyzed algal networks. We chose to manually add this reaction to the GSMNs of all five brown algae.

Pathways PWY-5915 and PWY-5917 generate phycobilins. As shown in Fig. 5, the *pebA* gene is associated with 1.3.7.2-RXN, *pebB* with 1.3.7.3-RXN, *pcyA* with 1.3.7.5-RXN, and *HY2* gene is with 1.3.7.4-RXN. In red algae, cryptophytes, and cyanobacteria, phycobilins bind to light-harvesting proteins called phycobiliproteins. Brown algae are thought to have lost phycobiliproteins during their evolution (Bhattacharya *et al*, 2004). However, studies suggested that brown algae have, in many cases, retained *pebA, pebB*, and *HY2* genes acquired via their secondary endosymbiosis event with red algae (Rockwell *et al*, 2014; Rockwell and Lagarias, 2020), that may be associated with the synthesis of phycobilins. The same authors propose that, in brown algae, expression levels of *pebB* could be low, and *pebA* sequences seem to have diverged between stramenopiles and other algal families (Rockwell and Lagarias, 2017). Globally the sequences predicted by AuCoMe for *E. siliculosus* also matched those identified manually in Cock et al (2010); Rockwell and Lagarias (2017), except that some *pebA* sequences were manually re-grouped in a separate clade named *pebZ* by Rockwell and Lagarias. However, according to AuCoMe, the *E. siliculosus pebA* and *pebB* genes were also associated with the reactions generally attributed to the *HY2* and *pcyA* genes. This is likely to constitute an artifact since these proteins share nearly 30% of sequence identity with *pebA* and *pebB*. The maintenance of phycobilin synthesis may be explained by the fact that these genes, although inherited from a red alga, have diversified and may now be used, e.g. to make new, unknown, photoreceptors rather than photosynthetic pigments (Rockwell and Lagarias, 2017). This example illustrates how complex it can be to interpret AuCoMe inferences in cases where conserved pathways provide the basis for potentially new functions in a given lineage.

### AuCoMe GSMNs are consistent with species phylogeny

To further assess the predictions of AuCoMe and to explore biological features, we clustered the GSMNs of all organisms of the algal dataset after the draft reconstruction as well as at the end of the pipeline by using the presence or absence of reactions in the GSMNs (see Fig. 6 A). The initial GSMNs produced from the annotations exhibited low consistency with the phylogenetic relationships among the algae (Strassert *et al*, 2021). Even well-established phylogenetic groups like red algae or brown algae were not recovered. At this step, the principal factor leading to the repartition of points in the MDS was the heterogeneity of genome annotations. An ANOSIM test supports this as it was not able to differentiate the main phylogenetic groups (R=0, P-value=0.45). However, in the MDS made from the GSMNs after the final step of AuCoMe, we observed a clear separation between the known phylogenetic groups, supported by an ANOSIM test (R=0.811, P-value=1e-04). This is also visible in the dendrograms clustering the GSMNs generated by the complete AuCoMe pipeline. This clustering was broadly consistent with the reference species phylogeny (Fig. 6B): there were only three higher order inconsistencies concerning *Cyanidiophora paradoxa*, for which the genome version deposited in Genbank was fully lacking expert annotations (Price *et al*, 2012), *Guillardia theta*, the phylogenetic position of which is controversial (Strassert *et al*, 2021), and *Nannochloropsis gaditana*, which was the only representative of eustigmatophycean stramenopiles. The two other stramenopile groups, diatoms and brown algae, were represented by multiple species which likely minimizes errors linked with peculiarities of a single genome. There were also some minor inconsistencies in intra-group relationships, in green algae, diatoms, brown algae, and opisthokonts.

**Figure 6:**
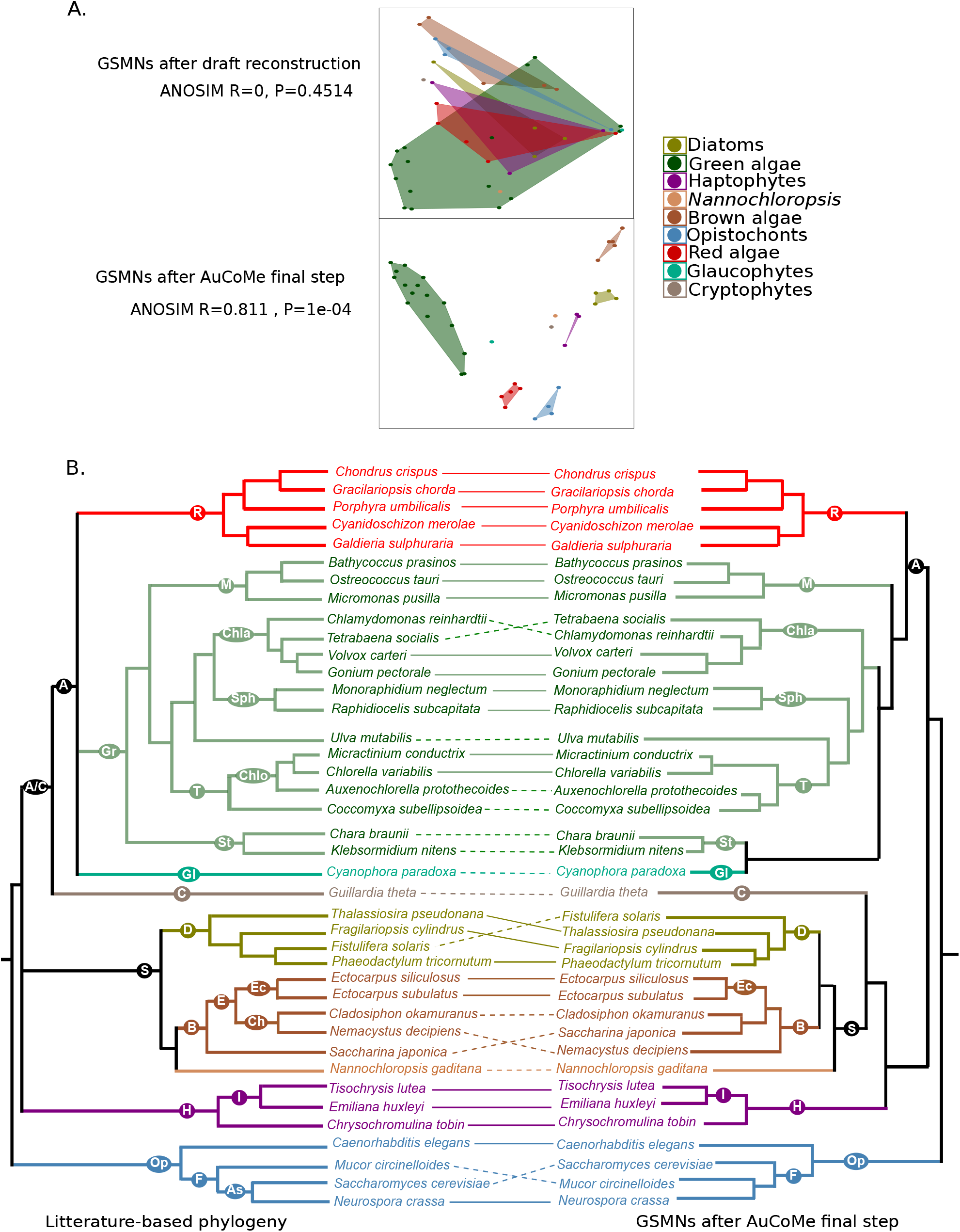
AuCoMe as a tool to improve taxonomic consistency of GSMNs. A. MDS plot for GSMNs calculated with the AuCoMe draft reconstruction step or after all AuCoMe steps. In both cases, ANOSIM values are indicated below (MDS and ANOSIM were computed using the vegan package (Oksanen *et al*, 2020)). B. Tanglegram evaluating the taxonomic consistency of AuCoMe dendrograms based on metabolic distances using the pvclust package (Suzuki and Shimodaira, 2006) with the Jaccard distance (right side) in comparison with reference phylogeny (left side), compiled from Strassert et al (2021). Full lines join species for which the position in the AuCoMe dendrogram is consistent with the reference phylogeny. Dotted lines join species for which the metabolic dendrogram and the reference phylogeny diverge. A/C: Archeplastids/Cryptophytes, A: Archeplastids, R: Rodophytes, Gr: Green algae, M: Mamiellales, Chla: Chlamydomonadales, Sph: Sphaeropleales, T: Trebouxiophyceae, Chlo: Chlorellaceae, St: Streptophytes, Gl: Glaucophytes, C: Cryptophytes, H: Haptophytes, I: Isochrysida, D: Diatoms, S: Stramenopiles, B: Brown algae, E: Ectocarpales, Ec: Ectocarpaceae, Ch: Chordariaceae, Op: Opistochonts, F: Fungi, As: Ascomycetes.

An illustration of the efficiency of AuCoMe was the *de novo* reconstruction of the GSMN of the glaucophyte *Cyanophora paradoxa*. For the reconstruction of this GSMN, we used the initially published genome sequence, which contained only two functionally-annotated genes (Price *et al*, 2012). The draft reconstruction by AuCoMe enabled us to retrieve 1,675 GPRs, a number within the same range as the other species from the dataset. Accordingly, *C. paradoxa* branched at the basis of the dendrogram after the draft reconstruction step, whereas it moved to the archeplastids after the orthology propagation step. Even if the grouping of *C. paradoxa* within archeplastids with the streptophytes *Chara brauni* and *Klebsormidium nitens* does not reflect the phylogenetic relationships, this shows that AuCoMe is a reasonable proxy for handling nearly unannotated genome sequences.

The small unicellular alga *G. theta* belonging to the cryptophytes grouped with stramenopiles rather than with the archeplastids or the haptophytes in our final GSMN dendrogram (Fig. 6B). Its plastid is derived from a secondary endosymbiosis event with a red alga (Curtis *et al*, 2012). Cryptophytes, together with haptophytes, have been proposed to form the supergroup of Hacrobia (Okamoto *et al*, 2009), and to belong to the kingdom of Chromista together with the stramenopiles (Cavalier-Smith, 2018). Alternatively, a common ancestor of cryptophytes and haptophytes may have been the source of the stramenopiles plastid via a tertiary endosymbiosis event (Archibald, 2009), and cryptophytes have also been suggested to be phylogenetically separate from haptophytes closer to the green algae lineage (Burki *et al*, 2012). To further examine the position of *G. theta* in our metabolic trees, we manually analyzed the presence/absence matrix of metabolic reactions to determine which of them most clearly linked *G. theta* to each of the three groups in question (stramenopiles, archeplastids, haptophytes). To this means we focused on reactions that distinguished at least two of these groups, i.e. that were present in at least 80% of the networks of at least one group, and absent from at least one other group (Appendix Table S7). A total of 216 reactions met this criterion, 109 of which were found in *G. theta* and 107 were absent. We then compared the network of *G. theta* to those of the other lineages regarding these reactions. We found that the network of *G. theta* shared the presence or absence of a similar number of distinctive reactions with all three groups: 120 with stramenopiles, 112 with haptophytes, and 101 with archaeplastids.

Next, we examined the metabolic pathways represented by the reactions that associated *G. theta* with the three groups, focusing on pathways that were ¿ 50% complete. The metabolic networks showed, for instance, that *G. theta*, (i) like haptophytes in our dataset, possess parts of the mitochondrial L-carnitine shuttle pathway, (ii) like the stramenopiles, comprises the complete pathway of glycine betaine synthesis, and, (iii) like terrestrial plants, can synthesize carnosine. We also manually examined the genes associated with these reactions, and found that in all cases, their sequences differed strongly from other sequences in the database, and could not be clearly associated with either archeplastids, stramenopiles, or haptophytes.

These examples underline the fact that cryptophytes diverged from the other lineages early in the history of eukaryotes and support the hypothesis that the metabolic capacities of extant cryptophytes might reflect adaptation to their specific environment more clearly than their ancient evolutionary history.

Another detailed analysis of the supervenn representation of the AuCoMe results (Appendix Fig S4) identified a cluster of six reactions common to the brown algae *Cladosiphon okamuranus* and *Saccharina japonica*, but surprisingly absent in other brown algal GSMNs (see Appendix Table S9). Four of these reactions correspond to putative enzymatic functions (glucuronate reductase, o-aminophenol oxidase, 11-oxo-*β*-amyrin 30-oxidase and keratan sulfotransferase), whereas the two others were predicted to occur spontaneously (RXN-13160 and RXN-17356). There was no obvious connection between these reactions as they are assigned to metabolic pathways involving different molecules. The presence of the four enzymatic reactions in *C. okamuranus* and *S. japonica* was assigned based on annotations, but orthology propagation in the AuCoMe pipeline identified only a subset of the potential orthologs (see Appendix Table S10). Manual analysis by blastP searches, however, showed that potential homologs of those four proteins were present in other brown algae in the dataset. This was further confirmed by the phylogenetic analysis of the o-aminophenol oxidases closely related to the *S. japonica* protein found by AuCoMe (SJ09941) and retrieved from the public databases (Appendix Table S10). Indeed, the phylogenetic tree showed that there are five clades of brown algal sequences present in three larger groups, including some stramenopile microalgae (see Appendix Fig. S5) Some more distant O-aminophenol oxidases found in *Symbiodinium* dinoflagellates and in fungi as well as metazoans form two well-separated branches from stramenopile proteins. These results suggest that the corresponding enzymes potentially exhibit other substrate or activity specificities in stramenopiles. The o-aminophenol oxidase family proteins present in the genome of *Ectocarpus siliculosus* are predicted to be cytoplasmic, extracellular, or to target the membrane (see Appendix Table S11), suggesting different roles depending on their subcellular localization. In this case, AuCoMe, with the support of more focused analyses, led to the identification of numerous candidate o-aminophenol oxidases in stramenopiles, whose biochemical functions and biological roles remain to be discovered.

## Discussion and conclusion

The numerous sequencing projects and available annotation approaches generate heterogeneously annotated data. There is currently a need to homogenize annotations across these data to make them comparable for wider scale studies. In this work we introduced a method to automatically homogenize functional predictions across heterogeneously-annotated genomes for large-scale metabolism comparisons between species across the tree of life. We illustrated how the tool can be applied both to prokaryotes and eukaryotes, even with high levels of annotation degradation.

### Accounting for existing annotations in the inference of homogenized GSMNs

Automatic inference of single species GSMNs is now routinely achieved, especially for prokaryotic species, and is often systematically performed for multiple genomes in order for instance to decipher putative functional interactions in microbial communities (Machado *et al*, 2018; Frioux et al, 2018). Such application requires a consistent quality of genomes and similar data treatment to be applied (genome annotation, metabolic network reconstruction) to minimize biases in predictions. More generally, one may aim at describing and comparing the predicted metabolism among related species from a given clade, and subsequently identify metabolic specificities. This endeavour is complex for eukaryotic genomes, which often need expert annotation as their enzymatic functions are much more difficult to characterize automatically. Moreover, annotation efforts can greatly vary between genomes, resulting in heterogeneous annotation and metabolic prediction quality.

The originality of our metabolic inference method resides in the possibility to account for, and preserve, available expert genome annotations. Not considering the genome annotations performed by specialists may lead to the omission of unique metabolic functions that are not well described in reference databases. On the other hand, comparing metabolic networks built from well-curated annotations to those built from poorly or automatically-annotated genomes will inevitably result in biases. In such cases, real metabolic differences between species cannot be distinguished from missing annotations in some genomes. AuCoMe constitutes a solution to such challenges through the propagation of expert annotations to less characterized genomes in the process of metabolic network reconstruction. By accounting for possibly missing functional but also structural annotations in the input genomes, the resulting metabolic networks are homogeneous and can therefore be directly compared.

### Method limitations and improvements

AuCoMe incorporates several strategies to optimize the method’s selectivity and sensitivity. Together these strategies collectively achieve comparable GSMN reconstruction, as demonstrated by validation experiments performed on randomly degraded *E. coli* genomes, although the results also highlight some of the method’s limitations.

A first limitation is illustrated by the comparison of AuCoMe reconstructions to the EcoCyc database considered as ground truth in our experiment. We observed that the GSMN automatically reconstructed from the reference genome substantially differs from the database. Extensive and systematic manual curation has been performed on this database since its creation in 1998 and we hypothesize that these efforts could not be all translated in the *E. coli* K–12 MG1655 annotations. As a result, several reactions were systematically missing from the automatic inferences provided by AuCoMe. This example illustrates the role of manual curation and expertise in producing high quality models. The homogenization of metabolic inference proposed by AuCoMe does not aim at replacing this step but rather enable an unbiased metabolic comparison between species.

Running AuCoMe on the bacterial dataset highlighted the impact of a single highly-annotated genome on metabolic inference. This dataset included the reference genome of the *E. coli* K–12 MG1655 strain which is better annotated than most of the other genomes considered. As a consequence, a certain number of reactions initially propagated by orthology from the *E. coli* K–12 MG1655 genome to others were discarded by the AuCoMe filter due to a lack of support, following the rationale that a single genome supporting an annotation propagation is not robust enough. Reasoning on ortholog clusters, the filter implies that several congruent genome sources are mandatory to confidently achieve an annotation propagation. While the relevance of the filter was demonstrated on the algal dataset by avoiding the propagation of annotations related to photosynthesis to non-photosynthetic organisms, it may be too stringent in some applications and lead to discarding relevant reactions. Several improvements of the filtering approach could be devised. For example, the structural annotation step could be improved: the annotation of pseudogenes in *Shigella* species would have been avoided by considering the annotations as pseudogenes available for the identified loci. More generally, in addition to the difficulties of automatically estimating protein homology, one must keep in mind that the link between orthology and conservation of function is still a matter of active investigation and methodological debate (Stamboulian *et al*, 2020; Begum et al, 2021).

Finally, we want to emphasize that our attempts to limit the inference of false positive reactions also directed the choice of method for the initial draft metabolic inference. We used Pathway Tools because of its several advantages such as the capacity to work with eukaryotic genomes, the suitability for parallel computing (Belcour *et al*, 2020a), and the possibility to limit gap-filling of metabolic networks. In addition, metabolic pathway completion performed by Pathway Tools does not systematically extend to ensuring the production of biomass. Pathway Tools was therefore adapted to our objective of avoiding to go beyond the strict interpretation of genome annotations. However, our perspective is to test other methods for the inference of draft GSMN from annotated genomes (Dias *et al*, 2015b; Zimmermann et al, 2021b), although some of them are restricted to prokaryotic genomes.

### Biological insights on the comparison of metabolic networks across species

The large-scale reconstruction and comparison of GSMNs made possible via AuCoMe will likely shed light on several fundamental biological processes, notably (1) the evolution of metabolic capacities, (2) the (metabolic) adaptations of organisms to their respective environments and modes of life, and (3) the metabolic interactions between symbiotic organisms.

### Evolution

Our examples of the Calvin cycle and phycobiliprotein synthesis demonstrate that, once all steps of the AuCoMe pipeline have been executed, the predicted metabolic capacities of the analyzed genomes reflect the biological knowledge we have of the corresponding organisms. Our approach, therefore, enables GSMNs to be compared in the light of evolutionary biology. The observation that metabolic dendrograms calculated from final AuCoMe reconstruction are mostly consistent with reference species phylogeny is in agreement with the literature. Indeed, numerous studies have shown that comparing GSMNs by computing a metabolic distance and clustering them into a dendrogram allow to cluster organism into groups close to the ones known by phylogenetic analysis but the position of species inside these groups is often different from the one of the phylogenetic groups (Vieira *et al*, 2011; Bauer et al, 2015; Prigent et al, 2018; Schulz and Almaas, 2020). It furthermore gives support to the hypothesis of a metabolic clock based on the congruence between molecular and metabolomic divergence in phytoplankton (Marcellin-Gros *et al*, 2020). The difference observed in the tanglegram 6 between phylogeny and metabolic distances could be further explored. One possibility could be to look at different similarity measures for the clustering. In this work, the Jaccard distance has been used but other measures could be used. For example, if we were to consider an absence of a reaction in two organisms as a similarity (to represent the loss of a function) then other measures could be envisaged such as the Simple Matching Coefficient. This also opens the perspective of inferring ancestral metabolic networks to better understand the dynamics of character evolution across time (Psomopoulos *et al*, 2020). One limitation of this approach, however, is that it relies on annotations already incorporated for at least one species in metabolic reaction databases, which are highly dependent on curation effort. Improving and systematizing the curation step will certainly become part of the standardization process necessary to gain full benefit of the ongoing massive sequencing effort at the global biosphere level (Lawniczak *et al*, 2022).

### Adaptation

The second aim of reconstructing comparable GSMNs is to determine to what extent metabolic changes are the result of or the prerequisite for adaptation. In our study, we made a first attempt at this question regarding the cryptophyte *G. theta*. This species has several potentially plesiomorphic metabolic traits in common with other marine lineages, that may constitute adaptations to their shared marine environment. Glycine-betaine, for instance, is known to be an osmoregulator or osmoprotectant in green plants (Di Martino *et al*, 2003), and carnosine has been proposed to function as an antioxidant in red algae (Tamura *et al*, 1998). Regarding carnitine, its physiological significance in photosynthetic organisms is still largely unknown, but antioxidant and osmolyte properties along with signaling functions have also been suggested (Jacques *et al*, 2018). However, for now, all of this remains purely hypothetical. To dig deeper into such questions in the future, we need to be able to distinguish changes that simply result from random processes such as metabolic drift (Belcour *et al*, 2020b) from changes that have an adaptive value. Currently, we envision two approaches that will help with this distinction. The first approach will be to further increase the number of species and lineages included in order to identify adaptive patterns, for example to among organisms occupying similar ecologcial niches. In phylogenomics, wide taxon sampling is recognized as one of the key features for reliable comparisons (e.g., (Young and Gillung, 2020)), whereas pairwise genomic comparisons across species are generally viewed as problematic (Dunn *et al*, 2018). Given that, as demonstrated above, phylogenetic signals in metabolism are stronger than the adaptive signals we can expect, this approach would also benefit from the development or adaptation of statistical models that could help detect signals of adaptation in an overall noisy dataset. Such models exist, for instance, to detect selective signatures in the evolution of protein-coding gene (Shapiro and Alm, 2008), but to our knowledge, have not been developed for metabolic networks or presence/absence signatures of genes. The second related strategy consists in focusing on phylogenetically closely related species that have only recently diverged and adapted to different environments. In such cases, we anticipate that the relative importance of drift along with the noise from the phylogenetic signal will be reduced due to the short evolutionary time since the separation. With such datasets, we may be able to reduce the level of replication required to find biologically relevant metabolic adaptations. The range of questions that could be addressed with the appropriate dataset is long and includes metabolic adaptations to different environments (Xu *et al*, 2020), food sources and domestication (Giannakou *et al*, 2020), multicellularity (Cock *et al*, 2010), or even life-history transitions to endophytism (Bernard *et al*, 2019).

### Interactions

Lastly, we anticipate that AuCoMe will provide new opportunities to study metabolic interactions between symbiotic organisms. For example, the tentative o-aminophenol oxidase activities pointed out by AuCoMe in brown algae could be involved in the protection against pathogen attacks at the cell surface. Indeed, a molecular oxygen-scavenging function in the chloroplast (Constabel *et al*, 1995) and a defense role (Gandía-Herrero *et al*, 2005) have been suggested for these enzymes in terrestrial plants. An o-Aminophenol Oxidase *Streptomyces griseusis* is known to be involved in the grixazone biosynthesis, i.e. an antibiotic (Suzuki *et al*, 2006). Similarly, brown algal o-Aminophenol Oxidases or Tyrosinases might be involved in the production of specific antibiotics. The o-Aminophenol Oxidase enzymes resemble laccases or tyrosinases. They can be involved in catechol or pigment production by oxidation (Jones *et al*, 1991). Numerous references have also shown that tyrosinases are efficiently inhibited by some phlorotannins, antioxidant compounds specific to the brown algae (Kang *et al*, 2004; Manandhar et al, 2019) suggesting there might be a regulation of polyphenol oxidation in certain conditions.

In the same vein, metabolic complementarity has previously been used to predict potentially beneficial metabolic interaction between a host and its associated microbiome (Frioux *et al*, 2018), and to successfully predict metabolic traits of the communities (Burgunter-Delamare *et al*, 2020). These studies have, so far, examined large numbers of symbionts (all sequenced and annotated with identical pipelines), but usually consider one specific host whose metabolic network was manually curated. With AuCoMe, these previous efforts could be expanded to incorporate a range of different hosts with their associated microbiota, thus facilitating the identification of common patterns in host-symbiont metabolic complementarity as well as their differences in these complementarities across different species and lineages. Just as for the question of adaptation, we believe this new scale of comparisons enabled by tools such as AuCoMe, will likely enable researchers to move from the study of specific examples to the identification of general trends, thus approaching the biologically most relevant evolutionary constraints.

## Methods

### Genomes and models

The *bacterial dataset* includes the 29 bacterial *Escherichia coli* and *Shigella* strains studied in (Vieira *et al*, 2011), downloaded from public databases (see Appendix Table S1 for details).

The *fungal dataset* includes 74 fungal genomes which were selected according to Wang *et al* (2009) as representative of the fungal diversity, together with 3 outgroup genomes: *C. elegans, D. melanogaster*, and *M. brevicollis*. All proteomes and genomes were downloaded from the NCBI Assembly Database (Kitts *et al*, 2016). The *algal dataset* contains 36 algal genomes selected to represent a wide diversity of photosynthetic eukaryotes and downloaded from public databases. The dataset includes 16 Viridiplantae (green algae), 5 Phaeophyceae (brown algae), 5 Rhodophyceae (red algae), 4 diatoms, 3 haptophytes, 1 cryptophyte (*Guillardia theta*), 1 Eustigmatophyceae (*Nannochloropsis gaditana*), 1 Glaucophyceae (*Cyanodophora paradoxa*). The genomes of *C. elegans* (Witting *et al*, 2018), *M. circinelloides* (Vongsangnak *et al*, 2016), *N. crassa* (Dreyfuss *et al*, 2013), and *S. cerevisiae* (Lu *et al*, 2019) were selected as outgroup genomes (see Appendix Table S3 for details).

Each annotated genome of the datasets was curated manually in order to make it compatible with Pathway Tools v23.5. Curated genomes are available at https://zenodo.org/record/6632441#.YqbMjS2FDOR.

### AuCoMe, a method to reconstruct genome-scale metabolic networks homogenized across related species

AuCoMe is a Python package implementing a pipeline whose steps are described in Fig. 1. The method aims at producing homogenized genome scale metabolic networks (GSMNs) for a set of heterogeneously-annotated genomes containing closely related or outlier species of a taxonomic group. AuCoMe takes as input GenBank files containing the genome sequences, the structural annotation of the genomes (gene and protein locations), the functional annotations (especially with GO terms and EC numbers) and the protein sequences (with the qualifier “translation” for the CDS feature). The output of AuCoMe is a set of GSMNs, provided in SBML and PADMET formats (Hucka *et al*, 2018; Aite et al, 2018). AuCoMe also produces a global report describing the sets of reactions added at all steps of the pipeline. The global panmetabolism, which is the complete family of metabolic reactions included in at least one GSMN of the set of genomes, is described in a tabulated file.

At **the initialization step** the command `aucome init` creates a template folder in which the user stores the input GenBank files.

The `aucome reconstruction` command runs **the draft reconstruction step**, which consists in reconstructing draft GSMNs according to the set of available genome annotations. During this step, the pipeline first checks the input GenBank files using Biopython (Cock *et al*, 2009). Then using the mpwt package (Belcour *et al*, 2020a), AuCoMe launches parallel processes of the PathoLogic algorithm of Pathway Tools (Karp *et al*, 2019). Pathway Tools creates Pathway/Genome Databases (PGDB) for all genomes. The resulting PGDBs are converted into PADMET and SBML files (Hucka *et al*, 2003, 2018) using the PADMet package (Aite *et al*, 2018). During this conversion, reactions from pathways with no enzyme identified in the genome (pathway hole reactions) predicted by Pathway Tools are removed as they are not associated with a gene and are not spontaneous reactions. For example, in Fig. 1 A, the draft reconstruction step generates 6 GPRs in total for the 3 considered genomes.

The `aucome orthology` command runs the **orthology propagation step**, which complements the previous GSMNs with GPRs associations whose genes are predicted to be orthologues to genes from GPR relations of other GSMNs of the dataset (Fig. 1 B). To that purpose, the pipeline relies on OrthoFinder (Emms and Kelly, 2015, 2019) for the inference of *orthologs* defined as clusters of homologous proteins shared across species. For each pair of orthologous genes shared between two species, the pipeline checks whether one of the genes is associated with an existing GPR association. In that case, a putative GPR association with the orthologous gene is added to the GSMN. At the end of the analysis of all genomes, a robustness score is calculated for assessing the confidence of each putative GPR association based on the number of annotated GPRs associations between the orthologs (see below). Non robust putative GPR associations are not integrated in the final GSMNs. In the example shown in Fig. 1 B, applying the robustness criteria leads to generating a putative new GPR association to the GSMN 2 (see the green orthogroup). In this example, the pipeline does not validate the GPR association related to the blue orthogroup because of insufficient annotation support.

The `aucome structural` command runs **the structural verification step** to identify GPRs associated with missing structural annotations of the input genomes. This pipeline step complements GSMNs with GPR associations from other GSMNs according to protein-against-genome alignment criteria. This enables the identification of reactions which are associated with gene sequences absent from the initial structural annotations of the input genomes. A pairwise comparison of the reactions in the GSMNs produced during the previous step is performed (Fig. 1 C). In this comparison, if a reaction is missing in an organism, a structural verification will be performed. For each protein sequence associated with a GPR relation in a GSMN, a tblastn (Altschul *et al*, 1990; Camacho et al, 2009) with Biopython (Cock *et al*, 2009) is performed against the other genome with the missing reaction. If a match (evalue¡1e-20) is found, the gene prediction tool Exonerate (Slater and Birney, 2005) is run on the region linked to the best match (region +-10 KB). If Exonerate finds a match, then the reaction associated with the protein sequence is assigned to the species where it was found to be missing. In Fig. 1 C, one reaction is added to the GSMN 2.

The command `aucome spontaneous` runs the **spontaneous completion step** to fill metabolic pathways with spontaneous reactions, in order to complement each GSMN obtained after the structural-completion step with spontaneous reactions, i.e. reactions that do not need enzymes to occur. For each pathway of the MetaCyc database (Caspi *et al*, 2020) which was incomplete in a GSMN, AuCoMe checks whether adding spontaneous reactions could complete the pathway. When this is the case, the spontaneous reaction is added to the GSMN. In Fig. 1 D, two spontaneous reactions are added to the GSMN 1 and GSMN 3. Then the final PADMET and SBML files are created for each studied organism.

### Availability and dependencies

AuCoMe is a python package under GPL-3.0 license, available through the Python Package Index at https://pypi.org/project/aucome. The source code and the complete documentation are available at https://github.com/AuReMe/aucome. Running AuCoMe on the datasets studied in the paper required as dependencies Blast v2.6.0 (Altschul *et al*, 1990), Diamond v0.9.35 (Buchfink *et al*, 2015), Exonerate v2.2.0 (Slater and Birney, 2005), FastME v2.1.15 (Lefort *et al*, 2015), MCL (Enright *et al*, 2002), MMseqs2 v11-e1a1c (Steinegger and Söding, 2017), OrthoFinder v2.3.3 (Emms and Kelly, 2015, 2019), Pathway Tools v23.5 (Karp *et al*, 2019). These following Python packages are needed to install AuCoMe: matplotlib mpwt v0.6.3 (Belcour *et al*, 2020a), padmet v5.0.1 (Aite *et al*, 2018), rpy2 v3.0.5, seaborn, supervenn, and tzlocal. The pvclust R package is also required. A docker or a singularity container can be created and enriched according to the dockerfile available on https://github.com/AuReMe/aucome/blob/master/recipes/Dockerfile.

### Robustness criteria for GPR association predicted by orthology

The robustness score of GPR associations of the pan-metabolic network was defined as illustrated in Algorithm 1 and detailed in the following. We denote by *org*(*g*) the organism of a gene *g*. For every pair of genes *g*1, *g*2 of two different organisms, we denote *orth*(*g*1, *g*2) = 1 if the genes are predicted to be orthologs. We denote by *association*(*r, g*) = 1 a GPR association between a reaction *r* and a gene *g* which is predicted by the AuCoMe algorithm. When the gene-association is predicted by the draft reconstruction step, we denote *annot type*(*r, g*) = 1 (and zero otherwise). When the gene-association is predicted according to orthology criteria, we denote *ortho type*(*r, g*) = 1 (and zero otherwise).

Let us consider now a reaction *r* of the pan-metabolic network. We denote by *N org*(*r*) the number of organisms for which the reaction *r* has been associated with a GPR relationship with any gene *g*: *N org*(*r*) = # {*org*(*g*), *association*(*r, g*) = 1} (L2, Alg. 1). For every gene *g* with *annot type*(*r, g*) = 1, we denote by *N prop*(*r, g*) the number of organisms different from *org*(*g*) the GPR association between *r* and *g* has been propagated to according to an orthology relation with the gene *g N prop*(*r, g*) = # *org*(*g*1), ∃*g*1 s.t. *org*(*g*1) ≠ *org*(*g*), *orth*(*g, g*1) = 1, *association*(*r, g*1) = 1. The GPR association between *r* and *g* is considered robust: *robust*(*r, g*) ≠ 1 as long as *annot type*(*r, g*) = 1.

The robustness assessment of a GPR between *r* and *g* propagated by orthology (L7, Alg. 1) distinguishes two scenarios. In the first scenario *g* belongs to an orthology cluster which is supported by at least two annotations. Formally this means that there exist two genes *g*1 to *g*2, both orthologs to *g*, such that *annot type*(*r, g*1)= 1 and *annot type*(*r, g*2)= 1. The presence of these genes leads us to consider *g* robustly associated with *r* (L8-9, Alg. 1).

In the second scenario the GPR association between *r* and *g* was propagated from a unique gene *g*1 with *annot type*(*r, g*1) = 1 in the orthology cluster (L11, Alg. 1). For these genes our strategy is to be as stringent as possible and we introduce a robustness criterion to reduce the risk of propagating false-positive reactions. The GPR association is considered robust if the number of organisms to which the reaction is propagated according to the annotation of *g*1 remains low with respect to the total number of considered organisms.

More precisely, *robust*(*r, g*) = 1 if *N prop*(*r, g*1) ≤ r*robust func*(*N org*(*r*) - 1) × (*N org*(*r*) - 1)l (L12-13, Alg. 1). The robustness function *robust* 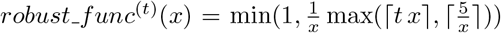 was chosen such that it is 1 for low values of *N org*, and then decreases to a threshold value (by default *t* = 0.05) for large values of *N org* (see a plot in Appendix Fig S1).

Altogether, the robustness criterion removes orthology predictions for GPR associations that are supported by a unique gene annotation and propagated to a large number of organisms. A toy example of the application of the algorithm is detailed in Appendix Section 2 and Fig S2).

#### Algorithm 1 Robustness criterion algorithm

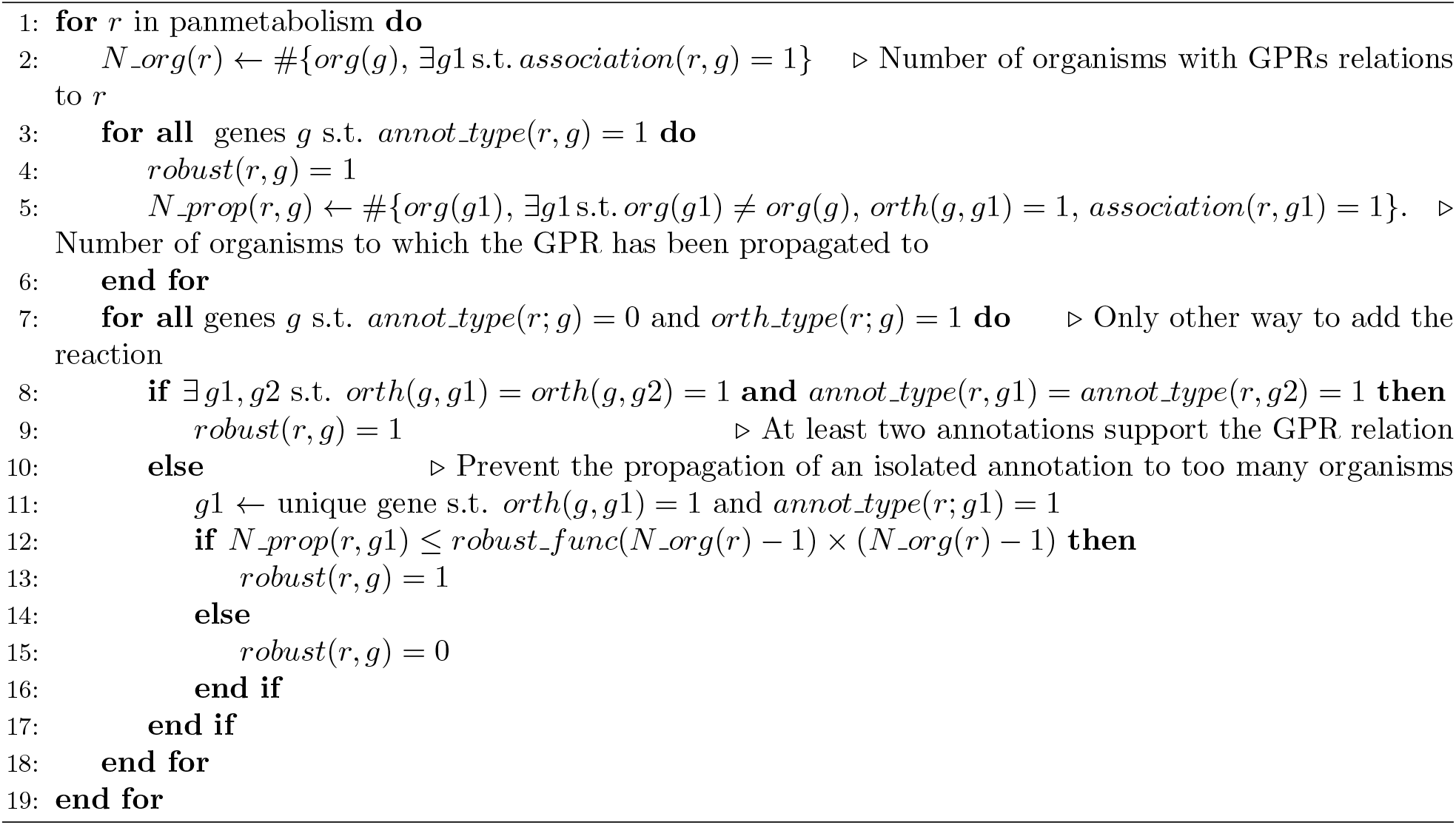

### Validation of the filtering step

One hundred random GPR associations were randomly selected and examined across the metabolic networks generated by AuCoMe for the algal dataset. Among them, 50 reactions that were predicted to be present and 50 reactions that were predicted to be absent in the metabolic networks. Regarding the former, their first associated gene was manually annotated based on reciprocal blast searches against Uniprot (Bateman *et al*, 2021) and the presence of conserved domains and the result of this manual annotation was compared to the predicted metabolic reaction. For absent reactions, we searched for characterized proteins known to catalyze the reaction in question, and then performed reciprocal BLASTp searches with the corresponding algal proteome.

In addition, DeepEC (version 0.4.0) (Ryu *et al*, 2019) was applied both to fungal and algal protein sequences. This tool predicts Enzyme Commission for protein sequences using 3 Convolutional Neural Networks trained on a dataset containing protein sequences associated with Enzyme Commission from SwissProt and TrEMBL. We extracted the EC numbers of reactions for which at least one GPR association was predicted according to orthology propagation for all reactions of the fungal and the algal datasets. For each EC number, we extracted the protein sequences associated with the considered reaction in the GSMNs, and we used DeepEC to infer an EC number for these proteins. Then we compared the EC number found by DeepEC (if found) to the EC number linked to the reaction by the pipeline.

### Comparison to the EcoCyc database

The complementarity between the orthology propagation step (second step) and the structural verification step (third step) was assessed using replicate genomes of *E. coli* K–12 MG1655 modified by a simulated degradation of its annotations. Degraded replicates were then given as input to the AuCoMe method, and the resulting GSMNs were finally compared to the ground truth Ecocyc metabolic network in order to estimate the precision and recall of the method. A non-degraded GSMN for *E. coli* K–12 MG1655 was obtained by running AuCoMe on its genome associated with 28 other *E coli* and *Shigella* genomes from Vieira et al (2011). This run allowed us to identify the genes associated with the metabolism that would be the target of the simulated degradation.

Then, the *E. coli* K–12 MG1655 genome was modified to generate replicates with randomly degraded annotations associated with GPR of the non degraded *E. coli* K–12 MG1655 GSMN. Two degradation types were simulated, (i) a degradation of the functional annotations of the genes, where all the annotations like GO Terms, EC numbers, gene names, etc. associated with a reaction were removed, and (ii) a degradation of the structural annotation of the genes, where gene positions and functional annotations were removed from the genome annotations. A third type of replicate was considered including the degradation of both structural and functional annotations. Replicates with increasing percentages of degraded annotations were generated for each of the three types of degradation. Furthermore the taxonomic ID associated with the *E. coli* K–12 MG1655 genome was degraded to *cellular organism*, to focus on the impact of genome annotations on GSMN reconstructions by AuCoMe, rather than on the effect of the automatic completion by the EcoCyc source performed by Pathway Tools when analyzing *E. coli* K–12 MG1655.

Each degraded replicate was associated with the 28 other *E. coli* and *Shigella* genomes, producing a synthetic dataset, which was analyzed by AuCoMe. This procedure therefore generated 31 synthetic bacterial datasets, plus the dataset with non-degraded *E. coli* K–12 MG1655 genome, which was called dataset 0. Their characteristics are detailed in Appendix Table S4.

For each *E. coli* K–12 MG1655 replicate in a dataset, AuCoMe produced a GSMN, which was compared to the metabolic database literature-based curated EcoCyc, considered as ground truth (Karp *et al*, 2002, 2018; Keseler et al, 2021).

We considered the reactions both present in a GSMN produced by AuCoMe and in the EcoCyc database as *True Positives (TPs). False Positives (FPs)* are reactions that are present in the GSMN produced by AuCoMe but not present in the EcoCyc metabolic network, and *False Negatives (FN)* are reactions present in the EcoCyc metabolic network but not present in the GSMN produced by AuCoMe. There were no True Negative reactions because each considered reaction either belongs to GSMNs produced by AuCoMe or the EcoCyc metabolic network.

The F-measure of each AuCoMe dataset was defined as 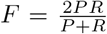 where 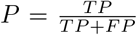 is the precision (number of reactions inferred by AuCoMe and present in EcoCyc among all the reactions predicted by AuCoMe) and 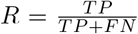 is the recall (number of reactions inferred by AuCoMe and present in EcoCyc among all reactions in EcoCyc).

### Phylogenetic analysis of the brown algal o-aminophenol oxidases

A dataset of 193 protein sequences was constructed using the closest homologues of the *S. japonica* o-aminophenol oxidase (SJ09941) in brown algae and extended to more distant sequences present in other organisms. Sequences were submitted to NGPhylogeny.fr via the A la carte” (Lemoine *et al*, 2019) pipeline as follows. The alignment was carried out by MAFFT (Katoh and Standley, 2013) using default parameters and automatically cleaned with trimAl (Capella-Gutiérrez *et al*, 2009) to obtain 372 informative positions. Then a maximum likelihood phylogenetic reconstruction was carried out using default parameters of the PhyML-SMS tool (Guindon *et al*, 2010; Lefort et al, 2017) allowing the best substitution model selection. Bootstrap analysis (Lemoine *et al*, 2018) with 100 replicates (¿ 70 %) was used to provide estimates for the phylogenetic tree topology. The Newick file (Junier and Zdobnov, 2010) was further formatted by MEGA v10.1.1 (Tamura *et al*, 2021) to obtain the simplified dendrogram (see Fig. 6(B)).

## Acknowledgements

We acknowledge the GenOuest bioinformatics core facility https://www.genouest.org for providing the computing infrastructure. We also acknowledge Pauline Hamon-Giraud for fruitful discussions. This work benefited from the support of the French Government via the National Research Agency investment expenditure program IDEALG (ANR-10-BTBR-04) and from Région Bretagne via the grant *<*SAD 2016 - METALG (9673)*>*.

## Supplementary Material

**Appendix File** The appendix file contains the description of the datasets, methodological details on the robustness criteria applied to a toy example, and additional details on the results on running times of the AuCoMe pipeline, validation of filtering steps and GPR associations, validation of EC numbers with deep-learning approaches, and analyses related to the consistency between AuCoMe GSMNs and species phylogeny.

**Additional file** The associated archive contains analyses (all tabulated files used to create the figures and results of the paper), datasets on which AuCoMe was run: the bacterial, fungal, and algal datasets, and the 32 synthetic datasets, which contain an *E. coli* K–12 MG1655 genome to which various degradations were applied, together with 28 other bacterial genomes. It is available at https://zenodo.org/record/6632441#.YqbMjS2FDOR.

## Author Contributions

**Arnaud Belcour**: Conceptualization, Data curation, Methodology, Formal Analysis, Software, Validation, Visualization, Writing – original draft, Writing – review & editing. **Jeanne Got**: Data curation, Formal Analysis, Resources, Software, Validation, Visualization, Writing – original draft, Writing – review & editing. **Méziane Aite**: Conceptualization, Data curation, Methodology, Software. **Ludovic Delage**: Formal Analysis, Validation, Writing – original draft, Writing – review & editing. **Jonas Collen**: Formal Analysis, Validation, Writing – review & editing. **Clémence Frioux**: Methodology, Software, Visualization, Writing – original draft, Writing – review & editing. **Catherine Leblanc**: Funding acquisition, Writing – original draft, Writing – review & editing. **Simon Dittami**: Conceptualization, Data curation, Formal Analysis, Funding acquisition, Methodology, Validation, Writing – original draft, Writing – review & editing. **Samuel Blanquart**: Conceptualization, Methodology, Writing – original draft, Writing – review & editing. **Gabriel V. Markov**: Data curation, Formal Analysis, Methodology, Supervision, Validation, Visualization, Writing – original draft, Writing – review & editing. **Anne Siegel**: Conceptualization, Formal Analysis, Funding acquisition, Methodology, Supervision, Writing – original draft, Writing – review & editing.

## Conflict of interest

The authors declare no conflict of interest.

